# Anterior claustrum cells are responsive during behavior but not passive sensory stimulation

**DOI:** 10.1101/2021.03.23.436687

**Authors:** Douglas R Ollerenshaw, Julianne Davis, Ethan G McBride, Andrew Shelton, Christof Koch, Shawn R Olsen

## Abstract

The claustrum is uniquely positioned to communicate with almost all higher-order cortical areas through widespread and reciprocal anatomical projections, yet the *in vivo* functional properties of claustrum neurons are not well understood. Here we use microendoscope imaging in mice to measure activity in populations of genetically-labelled Gnb4+ claustrum neurons. We find that only a small fraction of cells in the anterior claustrum are responsive to visual or auditory stimuli when delivered under passive yet wakeful conditions. In contrast, during a visual behavioral task, the majority of cells in the anterior claustrum are strongly modulated, with separate and spatially intermingled cell populations showing either increases or decreases in activity relative to spontaneous levels. Our results suggest that the Gnb4+ cells in the anterior claustrum do not represent passively presented sensory stimuli; rather, these cells are strongly engaged during behavior associated with sensory-motor transformations.

## Introduction

The claustrum is a subcortical brain region that is highly interconnected with the neocortex via widespread connections that are both direct and reciprocal (Torgerson et al., 2015; Wang et al., 2017; Zingg et al., 2018). Anatomically, the claustrum is a thin structure positioned between the neocortex and basal ganglia and is elongated along the anterior-posterior axis of the brain. Although many cortical regions connect reciprocally with the claustrum, in the mouse the majority of input connections from the cortex are from frontal and medial regions, with primary sensory areas providing relatively little input and receiving relatively few direct projections from the claustrum (Jackson et al., 2020; Wang et al., 2017; Zingg et al., 2018). These claustro-cortical and cortico-claustral connections show a general anterior-posterior topography, with more anterior parts of the claustrum connecting to more anterior cortical regions and more posterior regions connecting posteriorly (Wang et al., 2017). Based on its distributed connectivity, the claustrum has been hypothesized to subserve varied functions including consciousness (Crick and Koch, 2005), attention (Goll et al., 2015; Mathur, 2014; Smith et al., 2019), cortical synchronization (J. Smythies et al., 2014; J. R. Smythies et al., 2014), and sleep (Renouard et al., 2015).

Early studies using microelectrode recordings from the claustrum in cats and non-human primates reported basic sensory responses. Cells in the visual zone of the cat claustrum have large receptive fields and orientation tuning (LeVay and Sherk, 1981; Sherk and LeVay, 1983, 1981a, 1981b). In addition, responses to limb and trunk stimulation have been measured in the posterior claustrum (Olson and Graybiel, 1980). Remedios et al. found that claustrum neurons in macaque monkeys responded to the onset of either salient visual or auditory stimuli but not both (Remedios et al., 2014, 2010).

More recent studies have associated the claustrum with many diverse functional and cognitive roles including sensory change detection (Remedios et al., 2014), top-down action control (White et al., 2020, 2018), behavioral impulse control (Liu et al., 2019), navigation and spatial location coding (Jankowski and O’Mara, 2015), slow wave sleep (Narikiyo et al., 2020; Norimoto et al., 2020), attentional set shifting (Fodoulian et al., 2019), distractor suppression (Atlan et al., 2018), associative learning (Reus-garcía et al., 2020) and contextual association of reward (Terem et al., 2020). The diversity of functional properties measured in the claustrum could reflect spatially separate domains or cell types with distinct inputs. Indeed, a recent study found that cingulate-projecting CLA neurons were more likely to receive input from frontal areas, whereas non-anterior cingulate-projecting CLA neurons receive input from and project to sensorimotor regions (Chia et al., 2020).

In the mouse, there are numerous anatomical-genetic methods for labeling and expressing effector proteins in claustrum neurons, including viral injection (AAV) (Wang et al., 2017; White et al., 2020), retro-AAV injection into CLA projection areas (Jackson et al., 2018; Marriott et al., 2020; Zingg et al., 2018), CLA-Cre (Narikiyo et al., 2020), Egr2-Cre (Atlan et al., 2018; Terem et al., 2020), Esr2-Cre (Wang, 2019), D1-Cre (Terem et al., 2020), and Vglut2-Cre (Fodoulian et al., 2019). This results in a multiplicity of definitions of “claustrum neurons” and remains a challenge for the field.

We focus here on the Gnb4-IRES2-CreERT2 line. Gnb4 is a marker gene that labels glutamatergic neurons in the claustrum, dorsal endopiriform nucleus, and deep layers of some lateral cortical areas (Smith et al., 2018; Wang et al., 2017). The axonal projections of these Gnb4+ claustrum neurons have been reconstructed in their entirety and project widely to ipsilateral and cortical regions (Peng et al., 2021). We focus on the anterior claustrum and seek to characterize whether Gnb4+ cells in this region have sensory and/or behavior-modulated activity.

Physiological recordings from the claustrum *in vivo* are challenging to make given its thin, elongated structure. In this study, we used microendoscope imaging (Ghosh et al., 2011) to monitor the activity of genetically-labeled claustrum neurons in awake mice. We compared the activity of anterior claustrum neurons during passive sensory stimulation to activity during a task requiring mice to detect changes in visual stimuli (Garrett et al., 2020; Groblewski et al., 2020). We find that Gnb4+ cells in the anterior claustrum rarely respond to passively delivered sensory stimuli. In contrast, a majority of cells have robust responses during the visual behavioral task.

## Results

### Imaging activity of anterior claustrum cells

We used a miniaturized endoscope to image sensory and behavior-related activity in neurons in the anterior claustrum. A 1 mm diameter lens was chronically implanted above the anterior portion of the claustrum (Fig. 1A-C). We expressed the genetically encoded calcium sensor GCaMP6 in claustrum neurons using the Gnb4-Cre driver line crossed to a GCaMP6s or GCaMP6f reporter line, or injected with a Cre-dependent virus into the anterior claustrum (Fig. 1C,D). *Post hoc* histology verified the location of the micro-endoscope lens implant. Overall, we measured the activity of 956 cells from 12 mice (79.7 +/-46.7 cells across mice, range of 10 to 144). We identified individual cells using an ICA-based segmentation algorithm (Mukamel et al., 2009) and extracted z-scored fluorescence timeseries from each cell (Fig. 1D,E). Claustrum neurons showed low baseline activity, interrupted by large activity transients (Fig. 1E).

**Figure 1.**
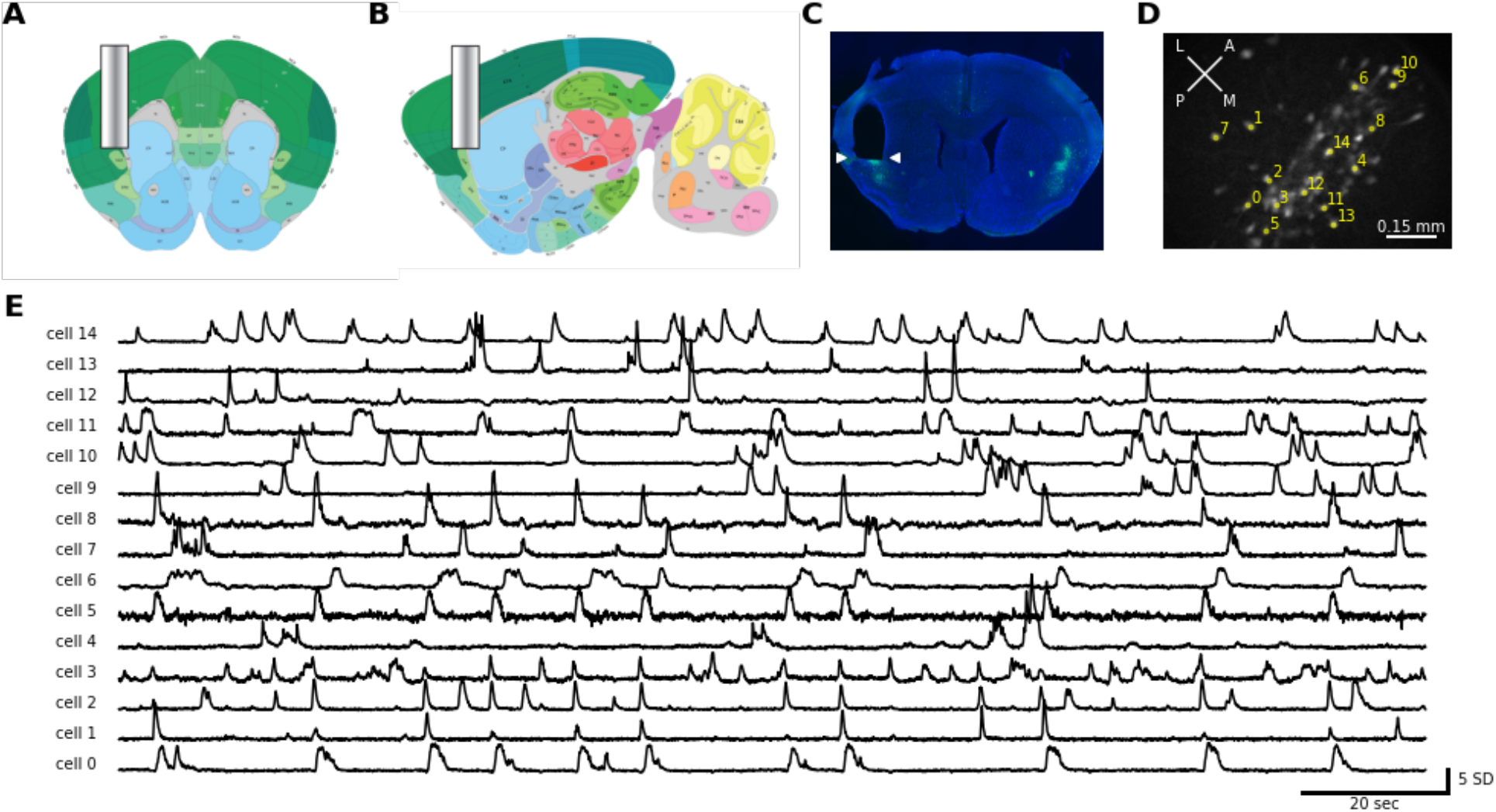
Microendoscope imaging of Gnb4+ claustrum cell populations. **A)** Schematic of coronal and **B)** sagittal plane showing approximate location of endoscope lens relative to anterior claustrum. Images taken from the Allen Mouse Brain Atlas. **C)** Histological section showing GCaMP6 labeled cells (green) and location of endoscope in one mouse brain. DAPI stain (labeling cell bodies) is shown in blue. Endoscope location is indicated by the white arrows and is visible as a void in the left hemisphere. **D)** Horizontal field of view in example mouse showing imaged Gnb4+ cells. Extracted ROIs are shown in white. Yellow dots and numbers correspond to timeseries shown in E. **E)** Spontaneous activity measured from 15 cells shown in D. Activity levels are expressed as a z-score.

In these chronically implanted mice, we measured Gnb4+ cell activity in multiple sessions spanning weeks. We first assessed activity before and during isoflurane-induced anesthesia. On a subsequent day, neural activity was monitored while sensory stimuli were passively presented to the mice. Finally, these same mice were trained on a behavioral task and we imaged activity during performance of this task.

### Anterior claustrum cells during anesthesia

We tested whether anterior claustrum cells were active during isoflurane anesthesia. We first made baseline recordings of ∼10 minutes while mice were awake in their home cages. Next, the animals were anesthetized in an acrylic induction chamber using 2-3% isoflurane, and subsequently transferred to an apparatus with a nosecone to continue isoflurane delivery while recordings were made at 2% then 1% isoflurane. Finally, mice were returned to their home cage to recover, and the session continued for ∼10 minutes after the animals became ambulatory. We recorded 676 cells from 8 animals in this experiment. Many cells were spontaneously active when the mice were awake in their home cage, but during 2% isoflurane anesthesia these cells were largely silent. Activity gradually increased after recovery from anesthesia, but often didn’t completely return to baseline levels during the 10-minute post-anesthesia recording (Figure 2A).

**Figure 2.**
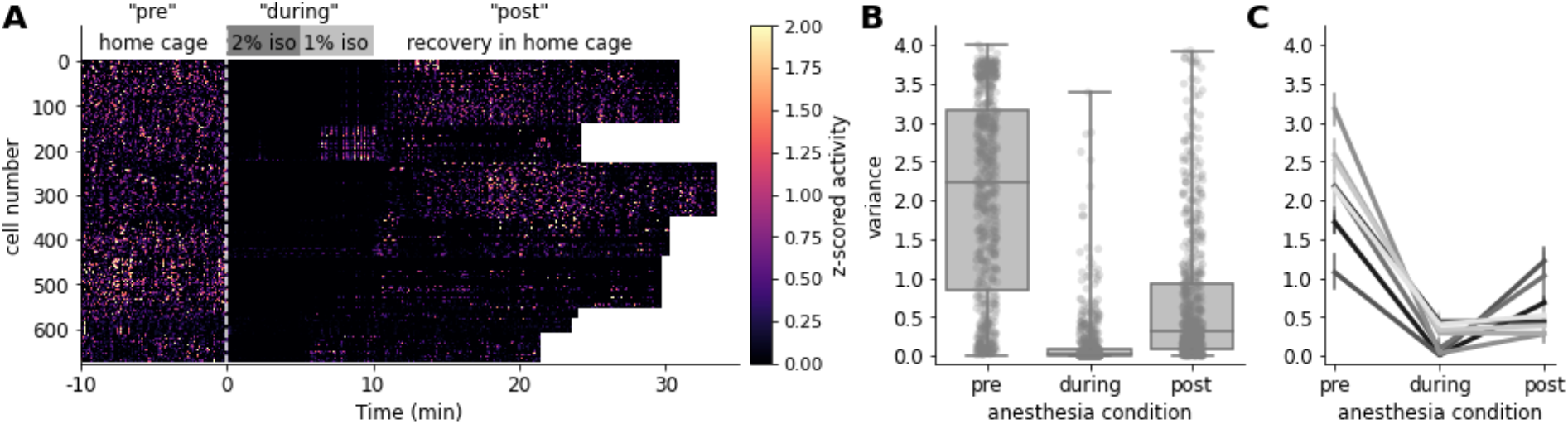
Claustrum cells are mostly silent during isoflurane anesthesia. **A)** Heatmap of z-scored fluorescence traces obtained from the claustrum in eight animals before, during, and after isoflurane anesthesia. Recordings are aligned to the time when recordings were restarted following isoflurane induction. Recording lengths varied across animals. **B)** Quantification of variance during three recording epochs: the pre-anesthesia period obtained in the home cage; the during-anesthesia period, when isoflurane anesthesia was being delivered via nosecone; and the post-anesthesia period, the first 10 minutes following return of the animal to its home cage. **C)** Same data as B, with each line representing one individual animal. Error bars represent 95% confidence intervals.

We quantified activity levels in this experiment as the variance of the z-scored fluorescence in one of three time bins: the home cage recording period, labeled “pre-anesthesia”; the 10 minutes of anesthesia at 1-2% isoflurane, labeled “during-anesthesia”; and the final 10 minutes of the recording in the home cage after the animal became ambulatory, labeled “post-anesthesia.” The variance of the z-scored activity dropped significantly from the pre-anesthesia to the during-anesthesia periods (pre-anesthesia mean = 2.07, median = 2.24; during anesthesia mean = 0.13, median = 0.012; p < 0.005), then increased in the post-anesthesia period (mean = 0.68, median = 0.32; p < 0.005). Note that activity in the post-anesthesia period remained significantly below that in the pre-anesthesia home-cage recordings (p < 0.005), indicating that longer than 10 minutes is required for activity in Gnb4+ neurons to return to baseline levels following isoflurane anesthesia.

### Passive sensory stimulation

We next examined whether anterior claustrum neurons were responsive to sensory stimuli presented to awake, head-fixed mice. On interleaved trials, we presented 200 ms long auditory (white noise) or visual stimuli (full field visual grating; Fig. 3A) with randomized, 3-5 s interstimulus intervals. For each cell, we quantified the trial-averaged activity within a 1.5 s window following the onset of the stimulus and compared that to activity in a 1.5 s baseline window preceding the stimulus to assess significant responses (see Methods). Few cells responded to visual stimulation (8/619, 1.3%), while only slightly more responded to the sound (22/619, 3.6%) (Fig. 3B-D). To test whether claustrum neurons responded to multimodal stimulation, we simultaneously presented both visual and auditory stimuli together on a subset of trials. A slightly higher fraction of cells (34/619, 5.5%) responded to the joint presentation of both stimuli; yet such multimodal responses remained rare (Fig. 3B-D). Of these 34 cells, just six responded on auditory-alone trials while only one cell responded on visual-alone trials. Together, these results indicate that cells in the anterior claustrum do not strongly encode passively presented sensory stimuli, contrasting with observations in more posterior regions of the claustrum measured in cats and macaque monkeys (Remedios et al., 2010; Sherk and LeVay, 1981a).

**Figure 3.**
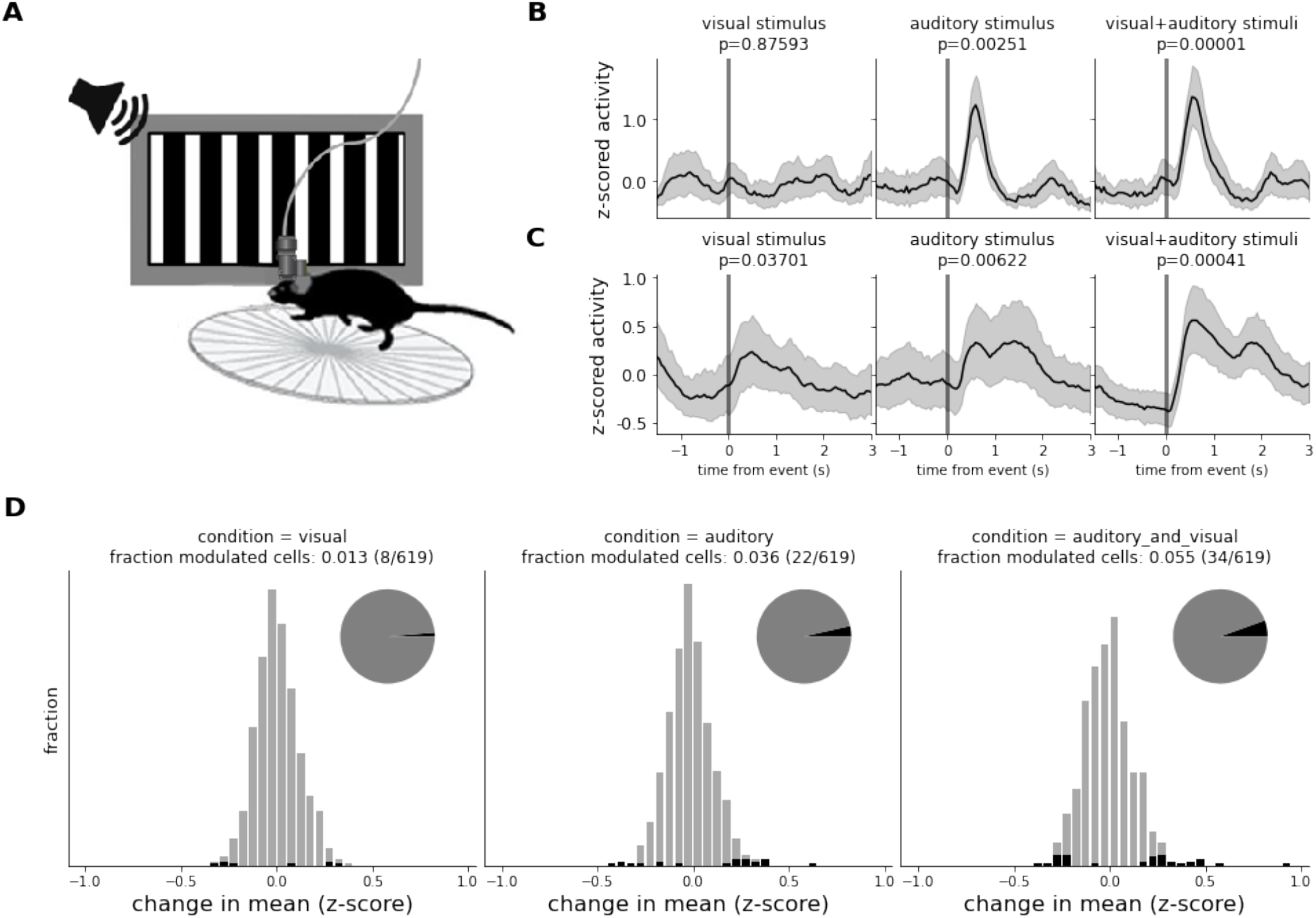
Few anterior claustrum cells respond to passively presented sensory stimuli. **A)** Schematic of passive sensory stimulation setup. Mice are head-fixed on a running wheel with a monitor to their right and a speaker nearby. Stimuli consist of full screen, 100% contrast square wave gratings and/or white noise bursts of 200 ms duration. **B)** Stimulus-locked activity in a rare cell that showed robust and significant responses to auditory and visual/auditory stimuli, but not to visual stimuli alone. **C)** A second example cell showing less robust, but statistically significant responses to all three stimulus conditions. **D)** Histograms of sensory responses magnitudes (measured as the mean response relative to baseline). Black bars represent significant responses at the p = 0.05 level and the inset pie charts show the fraction of all cells meeting this significance level.

### Activity during behavioral task

In the final experiment we tested whether anterior claustrum neurons were responsive during an active behavior, here a go/no-go visual change detection task (Fig. 4A, B) (Groblewski et al., 2020). Mice see a series of briefly presented (250 ms) images and are trained to respond to a new image by licking a spout. Water restriction is used to motivate mice to perform the task to earn water rewards. Mice were initially trained using high contrast visual grating stimuli before progressing to the natural scene images, requiring from 6-11 behavioral training sessions before reaching criterion performance. After training, we carried out microendoscope imaging of claustrum neurons in four behaving mice.

**Figure 4.**
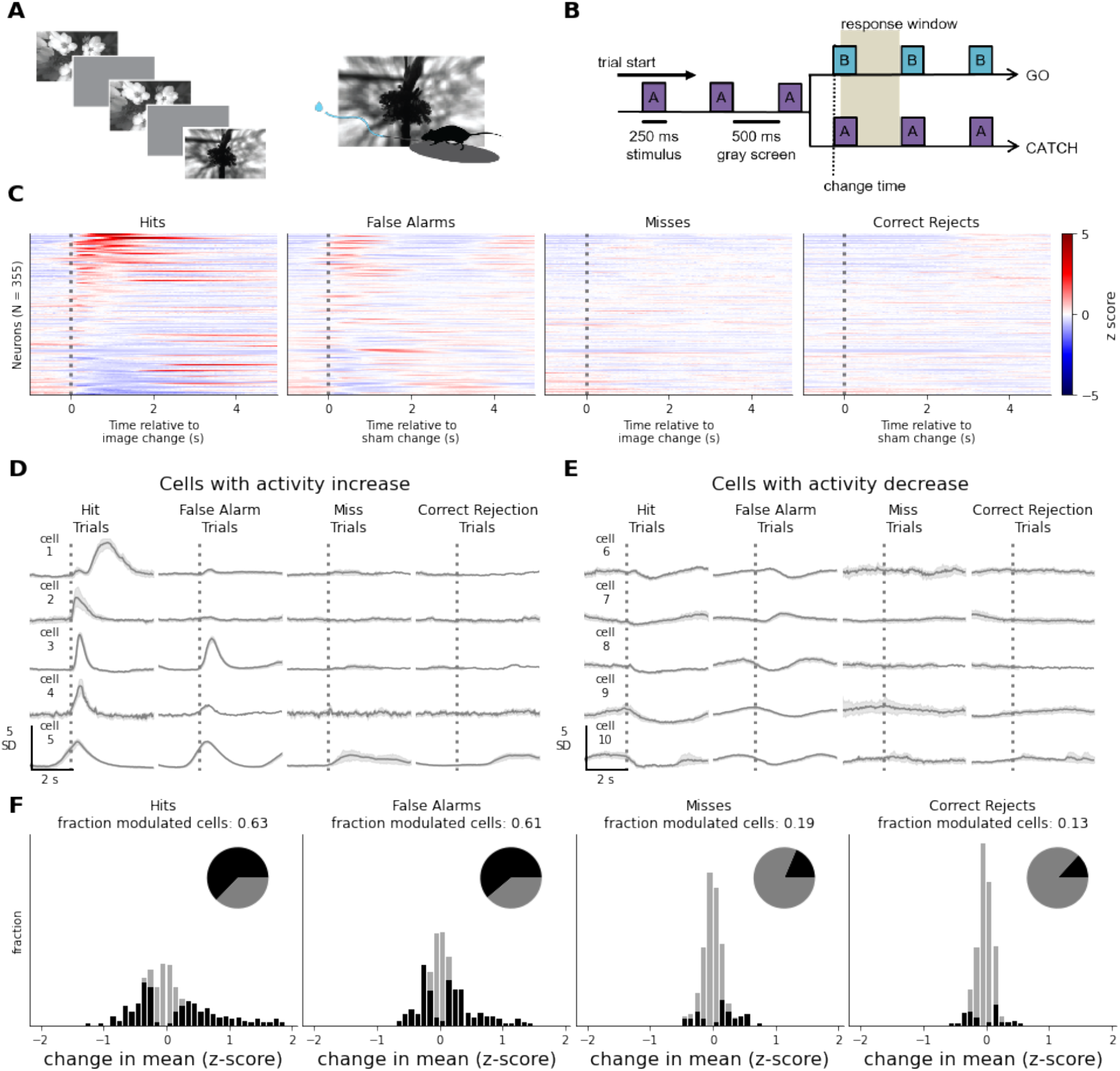
Claustrum cells show heterogenous responses during a visual task. **A)** Schematic of behavior. Head-fixed mice are trained to attend to a stream of repeatedly presented natural scenes and are rewarded for responding when a new scene is shown. **B)** On go-trials, the stimulus changes identity at a randomized time and mice are rewarded for licking in the response window following the change. On catch-trials, the stimulus identity is maintained and animal licks are counted as false alarms. **C)** Heatmap of event-triggered average activity aligned to the time of image change (or sham image change) for all cells (n = 355). The first column shows responses on hit trials, ordered by the response magnitude, and the following three columns show the responses for the same cells (vertical order maintained) on false alarm, miss, and correct rejection trials. **D)** Five exemplar cells that have significant activity increases on hit trials. Error bars represent 95% confidence intervals. **E)** Same as D for five different exemplar cells showing significant decreases in activity following hit responses. **F)** Histograms of response magnitudes for all recorded cells. Black bars represent significant responses at the p=0.05 level and inset pie charts show the fraction of all cells (in black) meeting this significance level. Over half of all cells show a significant response on hit or false alarm trials, in contrast to less than ∼3-5% under passive stimulus conditions (Fig. 3).

To analyze claustrum neuron responses, we separated trials according to trial-type: hits (correct response to a stimulus change); misses (stimulus change but no response); false alarms (response but no stimulus change); and correct rejections (no response following two consecutive presentations of the same image).

Many claustrum neurons showed substantial changes in activity around the time of behavioral responses on hit and false alarm trials (Fig. 4C,D). Overall, more than half of the anterior claustrum cells had significant activity changes on hit trials (223 of 355, 62.8%). A similar fraction of cells (217 of 355, 61.2%) showed significant modulation on false alarm trials in which the mouse made a behavioral response, but the stimulus didn’t change (Fig. 4E). Cells responsive on hit and false alarm trials were a largely overlapping population, with 55.6% of cells responding during hits or a false alarms also responding to the opposite trial type. Interestingly, different cells showed either increased or decreased activity relative to spontaneous baseline levels (Fig. 4D,E). Of the 223 significantly modulated cells on hit trials, 124 (56%) increased and 99 (44%) decreased their activity.

In contrast to hit and false alarm trials, most cells did not have a significant activity change on miss (18.6% responsive) or correct rejection trials (13.2% responsive) (Fig. 4F). These results indicate that many claustrum cells are engaged during this behavioral task and that these responses are most abundant when the animal makes a behavioral response (correctly or not).

On both hit and false alarm trials, the mouse decides to execute a motor action; thus, both motor planning and action could contribute to the claustrum activity we observe. Although a similar fraction of cells responded on both hit and false alarm trials, the magnitude of the cellular responses was significantly larger on hit trials (223 hit-responsive cells with a mean response magnitude of 0.64 standard deviations; 217 false-alarm responsive cells with a mean response magnitude of 0.37 standard deviations; p = 1.25e-11, independent two-tailed t-test). This suggests that additional factors beyond motor planning and action contributed to these responses. Whereas the behavioral response is the same on both hit and false alarm trials, only hit trials include both a stimulus change and a water reward for the animal. The larger responses on hit trials could be driven by the reinforced sensory stimulus change, in addition to the expectation and/or receipt of reward.

### Activated and suppressed cells are spatially intermingled

In our behavioral experiments, some cells showed strong increases in activity on hit trials while others decreased their activity. This suggests the existence of subsets of cells with divergent coding properties.

We next sought to evaluate whether cells with these different activity profiles are clustered within the imaging field of view. We visualized the spatial organization of cells with increased, decreased, or non-significant responses and found that these distinct groups are intermingled within the field of view (Fig. 5A). We quantified this observation by calculating the Euclidean distance between cell centers for every simultaneously recorded cell pair. If cells with the same response direction (positively or negatively modulated) would be clustered in space, the within-group distances should be less than the across-group distances, on average. In contrast to this prediction, we found that the distribution of distances for cells modulated in the same direction (orange) and opposite directions (blue) were completely overlapping, demonstrating that the within-group distances are not systematically lower than across-group distances (p=0.95, one-sided Mann-Whitney U test). This indicates that claustrum cells with different coding properties are intermingled within the same local region of the claustrum.

**Figure 5.**
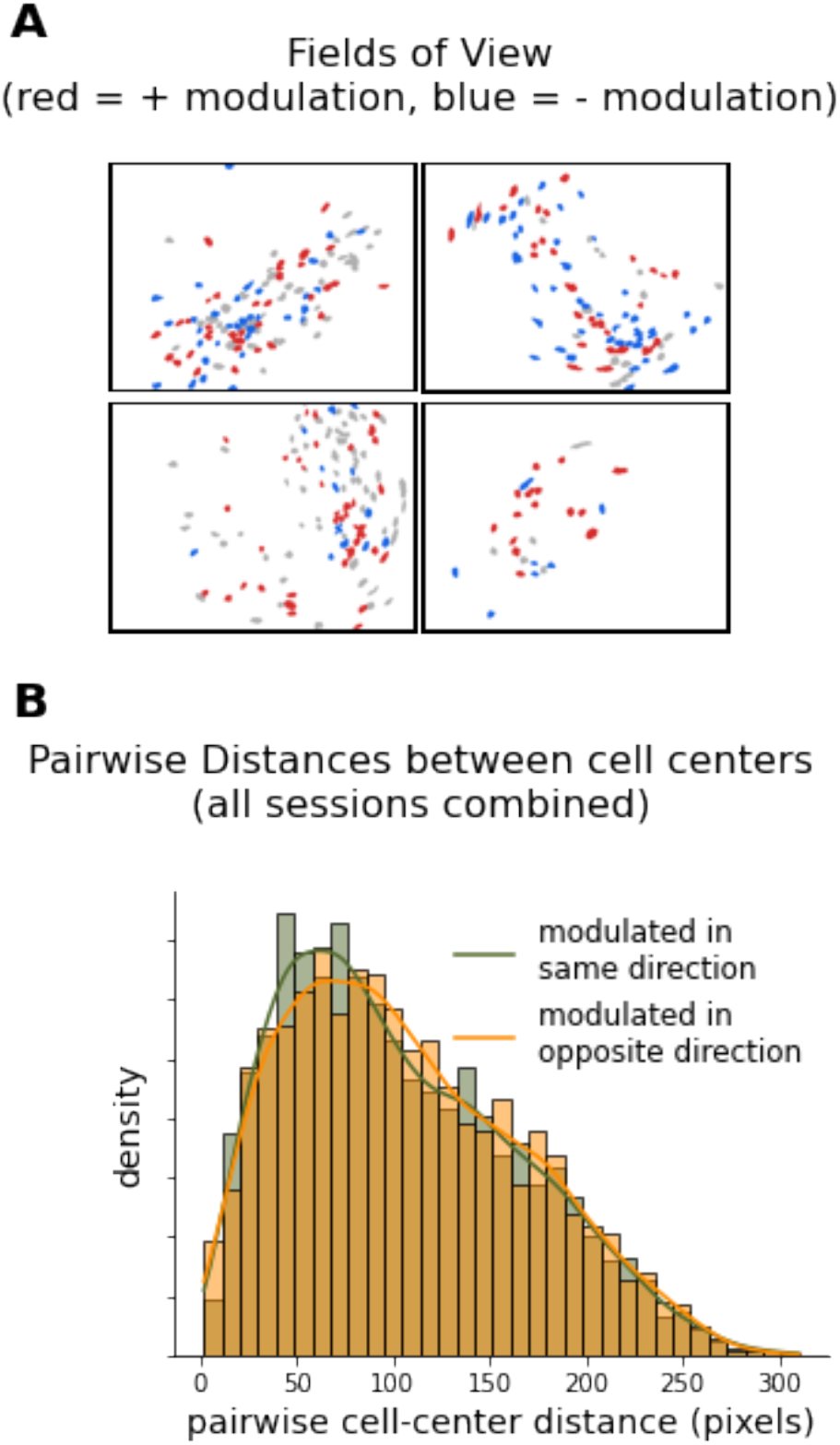
Claustrum cells with divergent responses are spatially intermingled. **A)** Fields of view from the four mice imaged during active behavior. The color of the cell ROIs indicate significant positive (red), significant negative (blue), or nonsignificant responses (grey). **B)** Euclidean distances between all simultaneously recorded pairs of cells with significant responses. Cells with the same or opposite modulation direction are equally distant from each other.

## Discussion

The conditions under which claustrum neurons are engaged *in vivo* are not well understood. Here, we found that activity of Gnb4+ anterior claustrum neurons is strongly modulated during a visual behavioral task, but not when visual or auditory stimuli are passively presented to mice; moreover, claustrum cells are virtually silent during isoflurane anesthesia. Our results imply the anterior claustrum is involved in processing during behavioral and cognitive tasks but is not engaged during passive sensory representation nor during anesthesia.

We used the Gnb4-Cre driver line to label claustrum neurons (Peng et al., 2021; Wang et al., 2017). It remains to be determined whether these Gnb4-positive cells represent a distinct functional group of cells within the claustrum, and whether Gnb4-negative cells have different activity patterns and functional roles. Also, the Gnb4-Cre driver line is not completely selective for the claustrum as some cortical layer 6 neurons as well as cells in the endopiriform nucleus are labeled (Peng et al., 2021; Smith et al., 2018; Wang et al., 2017). We collected data from Gnb4+ neurons by targeting the microendoscope lens implant to the claustrum and used *post hoc* histology for validation where possible. We verified that layer 6 cortical neurons were not in the field of view of our implants. However, since the endopiriform is directly ventral to the claustrum, we cannot completely rule out that some cells we imaged belonged to this structure. The relationship between the endopiriform and claustrum is a subject of ongoing debate (Smith et al., 2018).

In our behavioral task, we observed that many cells were significantly modulated during both hit and false alarm trials. This suggests the anterior claustrum could be involved in behavioral choices and motor responses, since both trial types include a decision and action by the mouse. Recordings in the primate claustrum have also reported cells with activity that precedes movements (Shima et al., 1996). However, in our study, response magnitudes were larger on hit compared to false alarm trials. Thus, perception of a stimulus change likely also contributes to claustrum activity. Consistent with this, on miss trials, when the mouse did not detect the stimulus change, responses were restricted to fewer cells and response magnitudes were smaller. Sensory change detection has been suggested as a function of the primate claustrum (Remedios et al., 2014). Overall, our data indicate that behaviorally relevant sensory, decision, motor and reward expectation components of the task could all contribute to claustrum cell responses. Future work should separately investigate how these variables shape claustrum cell activity during behavior.

We observed both enhancement and suppression of claustrum activity during the behavioral task. This indicates that bi-directional changes in claustrum activity could be important for communication with the cortex. The fraction of positively versus negatively modulated cells was roughly equal (35.1% activated, 27.8% suppressed, 37.1% not modulated). This bi-directional modulation of distinct groups of Gnb4+ claustrum cells could reflect competitive interactions among these. Although Gnb4-positive claustrum cells are glutamatergic and make excitatory synapses, the claustrum also contains GABAergic inhibitory cells that connect within the claustrum (Kim et al., 2016). These cells could mediate suppressive interactions between subpopulations of excitatory claustrum cells.

Recent studies in the mouse have shown that optogenetically stimulating the claustrum leads to suppression of pyramidal neurons in the cortex by activating cortical interneurons (Atlan et al., 2018; Jackson et al., 2018; Narikiyo et al., 2020). Thus, the increases in claustrum activity that we observe might serve to suppress specific cortical circuits, whereas the reduced activity in other claustrum cells could have a disinhibitory effect on separate circuits. Together, these bi-directional responses of separate claustrum populations could alter the balance between competing cortical populations. Indeed, an emerging theme in recent mouse studies is the finding that the claustrum helps to control attentional processes associated with distractor suppression, particularly during conditions of high cognitive load (Atlan et al., 2018; White et al., 2020, 2018).

The claustrum receives convergent input from many cortical areas and makes widespread and divergent projections back to the cortex. It is a much smaller brain region compared to the cortex, with a volumetric ratio of ∼1/200 in the mouse (Wang et al., 2017). This large difference in volume indicates the claustrum does not have the information coding capacity that the neocortex does. Thus, claustrum neurons may be more involved in coordinating the activity of the target cortical areas. Indeed, the axonal arbor of single claustrum neurons can target up to 29 distinct cortical areas (Peng et al., 2021). Our study suggests that the anterior claustrum might play a key role in coordinating cortical activity specifically during sensory-motor transformations associated with behavior and cognition.

## Materials and Methods

### Mice

All experiments and procedures were performed in accordance with protocols approved by the Allen Institute Animal Care and Use Committee. 6 male and 6 female mice expressing GCaMP6f or GCaMP6s in cells under Gnb4-Cre (Wang, 2017) control were used in the study. Both a viral expression strategy (3 mice) and a transgenic breeding strategy (9 mice) were used to contain expression to the desired cell population. Transgenic mice were back-crossed to C57BL/6J for two or more generations. Table 1 lists all mice used in the study, along with the sex, genotype specifics, and GRIN lens stereotactic implantation location relative to bregma (described below). All mice were maintained on a reverse light cycle.

**Table 1:**
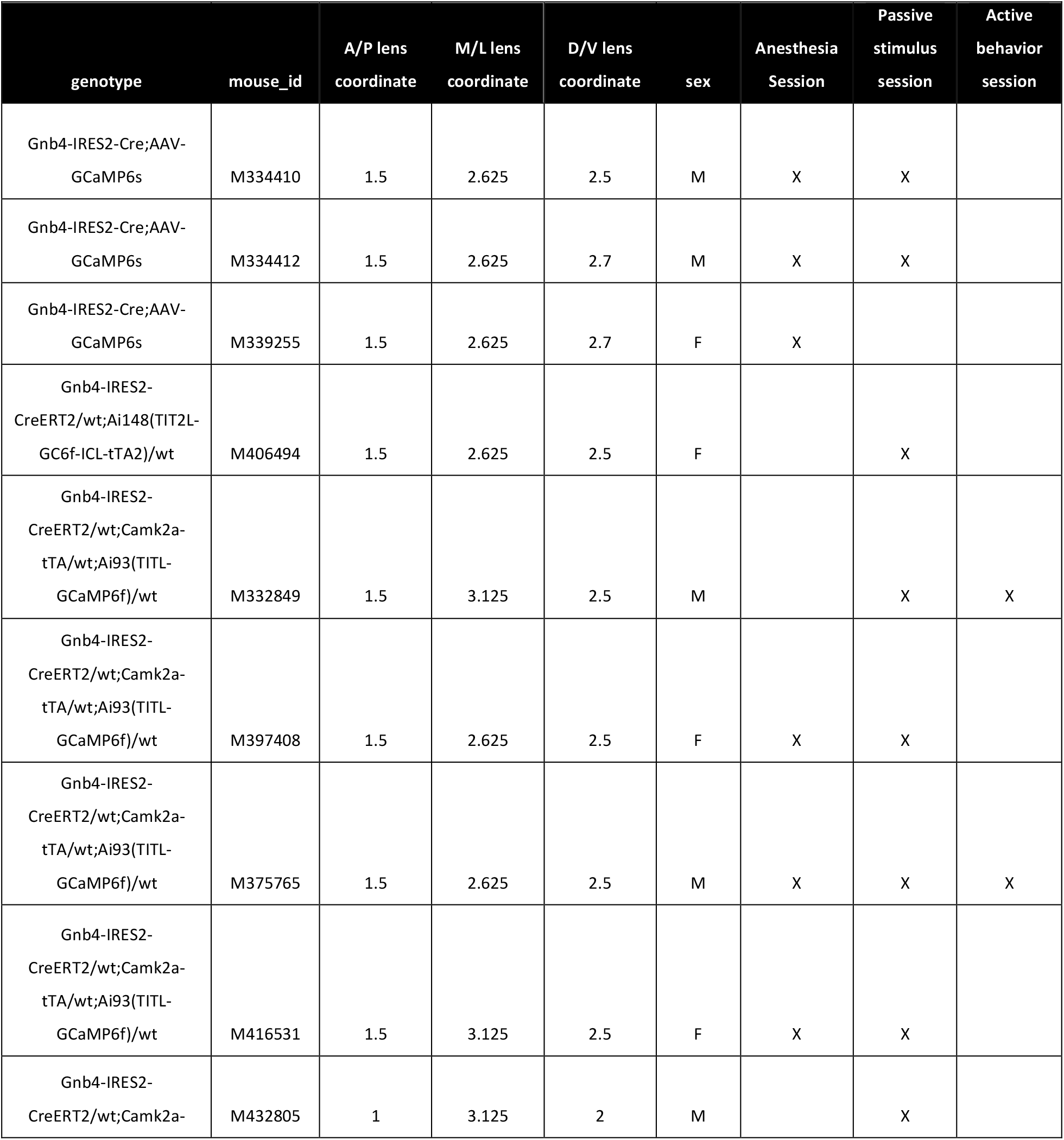
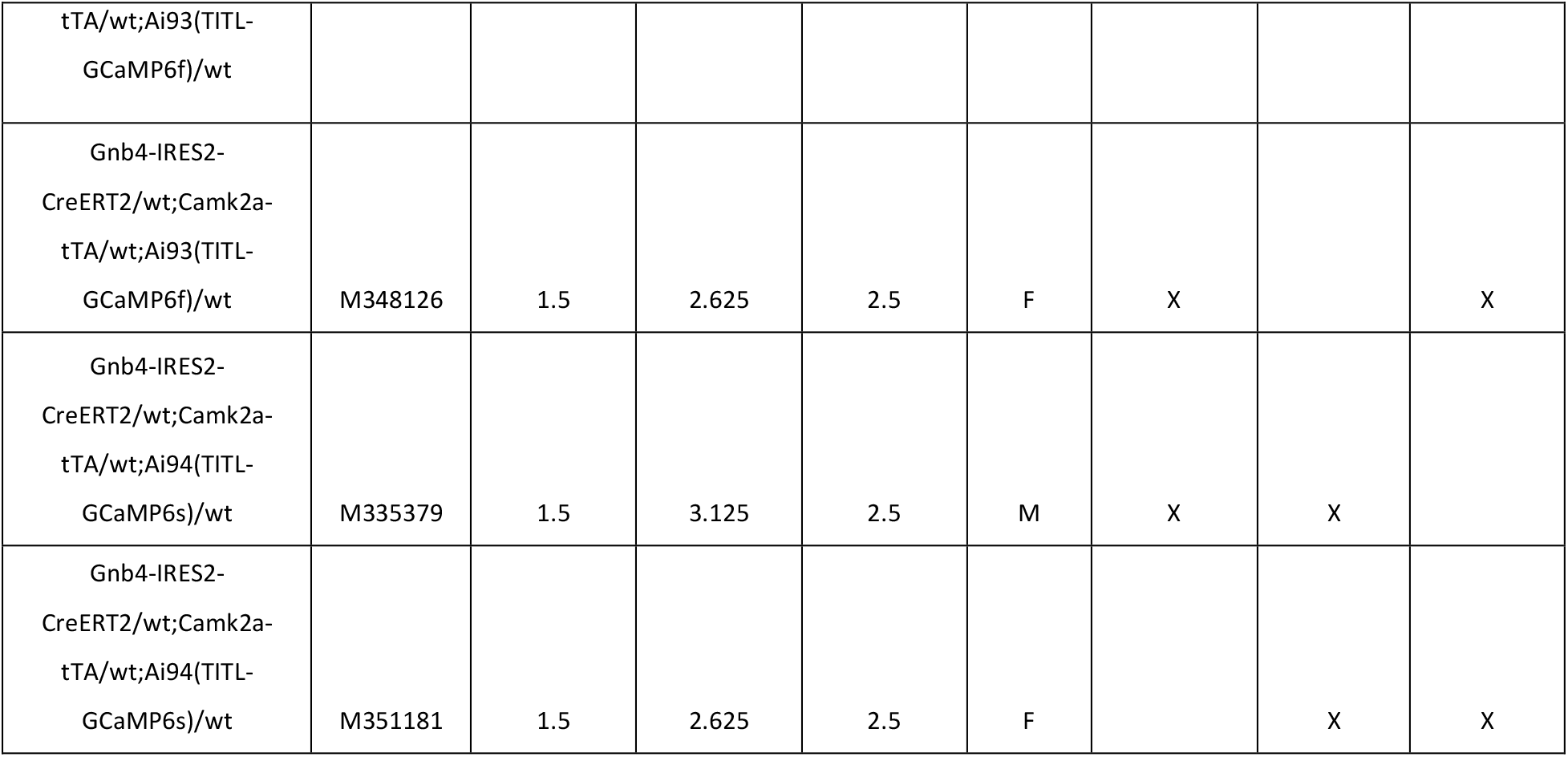
mice used in study with genotype, sex and GRIN lens implant coordinates listed

### Surgery

Surgery for GRIN lens and headpost implantation was performed on mice at least 5 weeks of age. All Gnb4-CreERT2 mice underwent 5 days of Tamoxifin induction via oral gavage (0.1 mL @ 50 mg/mL) prior to surgery, with at least 5 days of recovery following the last induction day prior to surgery.

1-3 hours prior to implanting the GRIN lens, mice were provided with an IM injection of Dexamethasone at 3-4 mg/kg and an SC injection of Ceftriaxone at 100-125 mg/kg. Mice were first anesthetized using 5% isoflurane in an induction chamber, after which they were transferred to a stereotaxic surgery rig (Kopf) and provided with 1.5-2% isoflurane via a nose cone. The eyes were covered with ophthalmic ointment for the duration of the surgery. Mice were provided with a pre-operative SC injection of Atropine (0.02-0.05 mg/kg) and Ketoprofen (2-5 mg/kg, respectively). Depth of anesthesia during surgery was monitored using breathing rate and toe pinch reflex. The surgical site was prepared by clipping then removing hair with depilatory cream, followed by topical application of betadine and 70% ethanol. The soft tissue overlying the skull was cleared and a 1-2 mm circular craniotomy was drilled at the desired stereotactic location over the left cortical surface, followed by removal of the dura with forceps. In the two mice receiving AAV injections, an injection pipette containing a GCaMP6s vector (AAV1.Syn.FLEX.GCaMP6s.WPRE.SV40) was slowly lowered to approximately 200 um ventral to the desired lens implantation depth, followed by a 2-5 minute wait to control for tissue deformation. 0.1-0.2 uL of viral solution was delivered via pressure injection at approximately 0.1-0.2 uL/minute, followed by a 10 minute wait, then a slow withdrawal of the injection pipette. For mice that did not receive an AAV injection, a guide path for the lens implant was created by slowly inserting a 20G needle to the desired insertion depth, then retracting.

1 mm diameter by 4 mm long Inscopix ProView GRIN lenses were then implanted by affixing the lens in the dedicated stereotactic guide clamp, then slowly lowering to the desired insertion depth. Fluorescence output from the lens was monitored using the Inscopix recording software during the implantation process, but cellular responses were never visible during the implantation process. After reaching the desired implantation depth, lenses were affixed to the skull using cyanoacrylate glue and metabond adhesive. A headcap was then built over the remainder of the exposed skull area using black metabond and a titanium headplate was secured with metabond over the posterior portion of the skull, approximately over the cerebellum. The top surface of the lens was covered in Kwik-cast for protection, then the animal was returned to a clean home cage on a heating pad for recovery. Post-operative Ceftriaxone and Ketoprofen was administered two times daily at the dosages listed above for the two days following implantation surgery.

In the weeks following surgery, and after at least 5 days of recovery, mice were periodically checked for GCaMP expression through the GRIN lens. To do this, mice were briefly head-restrained by clamping the head-plate in a custom clamp while the animal was free to run on a horizontal disc. The Kwik-cast plug protecting the GRIN lens was removed with forceps and the Inscopix microscope was clamped in a stereotactic guide and lowered over the dorsal end of the GRIN lens. If cells were not visible, the process was repeated 1-2 times per week for 4-6 weeks. If cells were visible, the microscope position relative to the lens surface was recorded, then the mouse was removed from head-fixation and briefly anesthetized to allow for installation of the Inscopix magnetic microscope baseplate. Isoflurane anesthesia was induced at 5% in an induction chamber, and the mouse was then transferred to a nosecone with isoflurane at 1-2% to maintain a steady, but light, level of anesthesia. The eyes were covered with ophthalmic ointment and the head was restrained using the implanted headbar. The microscope, with an attached baseplate, was lowered to the same approximate location relative to the lens surface at which cells were visible in the awake state. The baseplate was then cemented to the existing headcap using metabond, after which the microscope was removed, leaving behind the baseplate, and the animal was allowed to recover in preparation for subsequent imaging studies.

### Imaging studies

Imaging sessions were performed either while mice were in their home cage, in a nosecone under isoflurane anesthesia, or head-fixed and free to run on a rotating disc. Prior to the first head-fixed imaging session, mice were habituated to head fixation by gradually increasing the time of head fixation such that mice were able to sustain fixation for at least 30 minutes without overt signs of stress.

Passive stimulus sessions consisted of a 10-15 minute long imaging session in which mice were head-fixed and free to run on a behavior stage that was placed adjacent to a 1920x1200 pixel gamma-corrected LCD display at a viewing distance of 15 cm. The monitor was perpendicular to the right eye of the animal, thus limiting stimuli to a single monocular field. Custom Python code using the Psychopy2 library (Peirce et al., 2019) was used to control visual and auditory stimuli. Stimuli consisted of 0.04 cycle per degree full-contrast static square wave gratings (vertical and horizontal orientation randomly interleaved) presented for 200 ms and/or 200 ms white noise bursts delivered through standard PC speakers positioned near the animal. The three stimulus conditions (visual alone, auditory alone, or visual/auditory combined) were chosen with equal probability and inter-stimulus intervals were chosen on a uniform random distribution ranging from 3 to 5 seconds. Stimulus display times and microscope acquisition times were both fed as TTL pulses to a dedicated computer with a National Instruments PCIe board to allow for post-hoc time-alignment of signals.

Mice were subsequently placed on a water restriction protocol and trained in a go/no-go visual change detection task that has been described previously (Groblewski et al, 2020). Briefly, the task consisted of repeated presentations of images for 250 ms, separated by 500 ms of gray screen. Mice were rewarded with a small (7-10 uL) water reward for licking an adjacent lick spout in a brief (115 to 715 ms) window following the first presentation of a changed stimulus. Licks that preceded a change would result in a delay to the next change time, thus discouraging guessing. Training proceeded over series of stages with increasing task difficulty. Mice were initially trained on a variant of the task using static vertical and horizontal full-contrast grating stimuli. Stimuli were not interleaved with gray screens on the first training stage, resulting in sudden 90-degree shifts in the grating orientation at the change time. This was followed by a training stage in which the 500 ms gray screens separated each 250 ms grating presentation, with a 90 degree orientation change signaling the availability of reward. Finally, grating stimuli were replaced with natural image stimuli.

Change times were selected from a distribution with an exponentially decaying probability starting 2.25 seconds after the trial start. Licks that came before the change resulted in a short timeout, after which the delay to the next possible change was reset. On a subset of trials, denoted as catch trials, the image was held constant at the randomly selected change time, allowing for a direct measurement of chance performance. Therefore, after excluding trials in which the animal licked before the selected change time (referred to as ‘aborted trials’), there were four possible trial types: ‘hits’, or correct responses on go-trials; ‘misses’, or incorrect non-responses on go-trials; ‘false-alarms’, or incorrect responses on catch trials; and ‘correct-rejections’, or correct non-responses on catch trials. However, given that the false alarm count tended to be low in a given session, for the purposes of the analyses here, ‘false-alarms’ are defined as any response that was preceded by at least 2.5 seconds without another lick. Of the 12 mice that began the study, only four successfully learned the visual change detection task. In these four mice, we collected imaging data on a subset of sessions. To limit data volume and reduce the possibility of bleaching, data collection was limited to three non-contiguous 10-minute intervals within the full 60 minute behavior sessions. Time alignment between the acquired imaging frames and the visual stimuli was performed using the same TTL recording system described above.

### Data Analysis

All analyses were performed with custom routines written in Python 3.7.2 and open source packages. All processed data and the functions used to generate the figures in this study are publicly available at https://github.com/AllenInstitute/claustrum_imaging_manuscript.

Imaging data was processed using a processing pipeline built from the Inscopix Data Processing API (V1.1.2) for Python. The input to the pipeline, which is summarized briefly below, is a 20 Hz movie collected from the head-mounted microscope of green spectrum (500-550 nm) fluorescence emitted by GCaMP expressing cells in the GRIN lens field of view. These movies are spatially smoothed with a bandpass filter with a low frequency cutoff of 0.005 and a high cutoff of 0.5 to remove sensor noise, background, and other confounding signals.

The movies are then run through a motion-correction algorithm which identifies shared features across frames and translates images to compensate for motion. A key parameter of the motion correction algorithm is the maximum translation allowed on a given frame, which was set at 20 pixels.

Movies are then transformed to fractional changes in fluorescence, denoted DF/F, using the following formula:

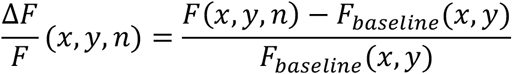

where *F*(*x, y, n*) represents the pixel value at a given spatial location *x, y* and frame number *n* and *F*_*baseline*_(*x, y*) represents the mean value of all movie frames at spatial location *x* and *y*.

The normalized movie is then run through the Inscopix PCA/ICA (principal and independent component analysis) algorithm to identify unique cell ROIs. This algorithm is based off of the published algorithm by Mukamel et al. (2009). Key inputs to the algorithm are the number of principal components to estimate, *PC =* 175; the number of independent components to estimate, *IC =* 1.15*∗ PC*; the unmixing type, which is set to ‘spatial’; the ICA temporal weight, which is set to 0; and the max iterations for ICA, which is set to 500.

The output of the PCA/ICA step is an array of shape *PC x N*, where *PC* is the number of estimated principal components and *N* is the number of frames, representing the estimated activity of each candidate cell at each point in time. This array is then passed to an event detection algorithm that attempts to automatically detect bursts of cell activity from the continuous PCA/ICA output. Key parameters of the event detection algorithm are the threshold, in median-absolute-deviations, that the trace has to cross to be considered an event, which was set to 5; the value *tau*, which is the minimum event duration, and was set to 300 ms; and the event time reference, defined as the event maxima, which specifies the time that will be assigned to the event. By default, negative-going transients are ignored in the event detection algorithm.

Finally, candidate cells from the PCA/ICA algorithm were passed through an automated accept/reject stage in which only cells with an SNR or 3 or greater, an event rate greater than 0, and a continuous spatial extent (i.e., the identified ROI did not consist of multiple, disconnected components across the field of view) were accepted. Additionally, potentially duplicated ROIs were identified as those with centers within 10 pixels, temporal correlations of 0.7 or greater, and structural similarities of 0.95 or greater (with structural similarity calculated using scikit-image.measure.compare_ssim). In cases where duplicate ROIs were found, the cell with the highest SNR value was retained. Finally, timeseries from the PCA/ICA algorithm were detrended by subtracting a 1000 sample (50 second) median-filtered version of the trace, then z-scored. All subsequent analyses were performed on the detrended and z-scored traces. The raw and detrended/z-scored timeseries are available in the github repository for every cell included in the study.

Activity from the imaging sessions was analyzed by calculating the mean of the z-scored timeseries in a 1.5 second window preceding each stimulus event and an equally sized 1.5 second window following every event. In the passive sessions, events were defined as the stimulus onset times for each of the three unique stimulus types: visual stimuli alone, auditory stimuli alone, or visual and auditory stimuli combined. In the active behavior sessions, events were defined as follows: hits – the time of the stimulus change on go-trials where the animal licked in the response window, defined as a 115 to 715 ms window following the display-lag-compensated image display time; misses – the time of the stimulus change on go-trials where the animal did not emit a lick in the response window following a change; correct-rejections – the time of a no-go stimulus on catch trials in which the animal correctly withheld response; false alarms – the stimulus display time preceding the first lick following a minimum of 2.25 seconds without a lick on trials without a stimulus change. Mean activity preceding all events was pooled to form a distribution of pre-stimulus values for each cell. Responsive cells were identified by comparing the distributions of activity preceding all events with that following each of the unique event types using a two-tailed independent t-test with an alpha level of 0.05.

Imaging during the 60 minute behavior sessions was limited to three 10 minute epochs, separated by two approximately 10 minute periods with the camera and excitation LED powered off. This was designed to avoid photo-bleaching and/or excessive heat buildup and was introduced after all cells disappeared from the field of view over a 2-3 day period in an early pilot mouse.

Imaging data from the three distinct recording epochs were combined using the Inscopix cell-matching algorithm. In one experiment, six ROIs that had passed other acceptance criteria were manually rejected due to inconsistencies in the timeseries after cell-matching.

The degree of spatial overlap of positively modulated vs negatively modulated cells during the behavioral task was quantified by identifying the location of the center of mass of each significantly modulated cell ROI. The Euclidean distance was then calculated between each pair of simultaneously recorded ROIs. The distribution of Euclidian distances for cell pairs that were modulated in the same direction (both positive or both negative) was compared to the distribution of Euclidean distances for cell pairs that responded in opposite directions (one positive and one negative) using the one-sided non-parametric Mann-Whitney U-test implanted in scipy (v1.2.0).

## Acknowledgements

We thank the Allen Institute founder, Paul G. Allen, for his vision, encouragement, and support. We thank the Tiny Blue Dot Foundation for providing funding for this study. We thank Jacqulyn R. Kuyat for assistance with histology. We thank Rylan Larsen for assistance with viral vectors and valuable input during experimental design. We thank Quanxin Wang and Rachel Tompa for useful feedback on the manuscript. We thank Chris Tsang and Kai-Siang Chen, field scientific consultants from Inscopix, for their assistance with methods development, experimental design, and analysis. We also thank Mike Schacter, software engineer from Inscopix, for assistance in setting up the data processing pipeline.

## Notes

https://github.com/AllenInstitute/claustrum_imaging_manuscript

### Competing Interest Statement

The authors have declared no competing interest.

